# Racing against stomatal attenuation: rapid CO_2_ response curves more reliably estimate photosynthetic capacity than steady state curves in a low conductance species

**DOI:** 10.1101/2020.08.28.270785

**Authors:** C Vincent, MO Pierre, JR Stinziano

## Abstract

*A/C*_i_ curves are an important gas-exchange-based approach to understanding the regulation of photosynthesis, describing the response of net CO_2_ assimilation (*A*) to leaf internal concentration of CO_2_ (*C*_i_). Low stomatal conductance species pose a challenge to the measurement of *A/C*_i_ curves by reducing the signal-to-noise ratio of gas exchange measures. Additionally, the stomatal attenuation effect of elevated ambient CO_2_ leads to further reduction of conductance and may lead to erroneous interpretation of high *C*_i_ responses of *A*. Rapid *A/C*_i_ response (RACiR) curves offer a potential practice to develop *A*/*C*_i_ curves faster than the stomatal closure response to elevated CO_2_. We used the moderately low conductance *Citrus* to compare traditional steady state (SS) *A*/*C*_i_ curves with RACiR curves. SS curves failed more often than RACiR curves. Overall parameter estimates were the same between SS and RACiR curves. When low stomatal conductance values were removed, triose-phosphate utilization (TPU) limitation estimates increased. Overall RACiR stomatal conductance values began and remained higher than SS values. Based on the comparable resulting parameter estimates, higher likelihood of success and reduced measurement time, we propose RACiR as a valuable tool to measure *A*/*C*_i_ responses in low conductance species.

## Introduction

Gas exchange is a powerful tool in plant physiology, allowing us to understand CO_2_ and water relations in plants to compare plant performance in a range of environments (e.g. Stinziano & Way, 2017; Vincent et al., 2017; Smith et al., 2020), to understand why and how plants cope with stressful environments (e.g. Kumarathunge et al., 2019; Zhu et al., 2020), and for modeling plant-atmosphere interactions up to the global scale (e.g. Oleson et al., 2013; Rogers et al. 2017; Lombardozzi et al., 2018). However, gas exchange measurements require a sufficient signal-to-noise ratio to obtain quality data for inferences. To obtain an adequate signal-to-noise ratio in a leaf, stomata need to be sufficiently open for a quality signal.

Challengingly, plant species present a wide variation in stomatal conductance depending on ecotype and environment, with many exhibiting low conductance, with maxima below 0.2 μmol m^−2^ s^−1^ (Körner et al. 1986; Tezara et al. 1998), and rapidly stomatal closure whenever conditions are not ideal (Radin et al., 1994) or at a particular time of day (Steppe et al., 2006). For example, in a study of *Citrus x sinensis* (L.) and *Citrus reticulata* (Blanco) in which treatment did not affect stomatal conductance, 35% of gas exchange measurements exhibited stomatal conductance to water (*g*_sw_) below 0.08 μmol m^−2^ s^−1^ (Vincent et al., *unpublished data*). When stomatal conductance is low, not only is the signal-to-noise ratio low in the gas exchange measurements, it becomes difficult to obtain a sufficient range in intercellular CO_2_ concentration (*C*_i_) to characterize the response of net CO_2_ assimilation (*A*) to *C*_i_ (*A/C*_i_ curves) (Stinziano et al., 2020). Further compounding the signal-to-noise challenge in the context of *A/C*_i_ curves, plants respond to elevated CO_2_ by stomatal closure on an order of minutes.

*A*/*C*_i_ curves are a powerful tool to which we can apply the Farquhar-von Caemmerer-Berry model of photosynthesis (Farquhar et al. 1980) to estimate photosynthetic biochemistry for comparing plant performance (e.g. Way & Sage, 2008), understanding photosynthetic acclimation (e.g. Kumarathunge et al. 2019), and modeling vegetative carbon uptake (e.g. Oleson et al., 2013). In species with low stomatal conductance or stomata that close rapidly at high CO_2_, these responses are very difficult to characterize. Stomatal limitations may further lead to incorrect conclusions on what is limiting photosynthesis. For example, a stomatal limitation may be incorrectly interpreted as a limitation due to rubisco carboxylation. The recent development of rapid *A/C*_i_ curves (RACiR, Stinziano et al. 2017, 2019ab), may be sufficiently fast to circumvent the difficulties of stomatal closure in such species.

Here we demonstrate the benefits of RACiR, relative to steady state *A/C*_i_ curves, in so-called “difficult” species using *Citrus x sinensis*, a species that is difficult to work with due to rapid stomatal fluctuations and moderately low stomatal conductance (Steppe et al. 2006; Syvertsen 1982).

## Materials & Methods

### Plant material

*Citrus sinensis* (L.), cultivar ‘Valencia,’ plants were grown in a greenhouse in Lake Alfred Florida (28.1021 °N, 81.7121 °W) in a natural diurnal cycle and daily maximum and minimum of ~25-34 °C. Plants were irrigated daily and fertilized according to standard practices. Steady-state and RACiR curves were performed on leaves of the same plants in a paired fashion on 19 plants for 19 RACiR curves and 19 steady-state (SS) curves.

### Gas exchange

Response curves were measured using an infrared gas analyzer (LI-6800, LiCor Inc., Lincoln, NE). Leaf chamber settings were fixed at 60% relative humidity, 1,200 μmol m^−2^ s^−1^ photosynthetically active radiation, and set to maintain the ambient temperature (31-34 °C). For all measurements we used a threshold of *g*_sw_ of 0.08 μmol m^−2^ s^−1^. If *g_sw_* was below this threshold before beginning, the response curve was not initiated. RACiR data were gathered at 0.5 Hz, and began by stabilizing measurements at a reference CO_2_ concentration of 100 μmol mol^−1^ at a ramping rate of 100 μmol mol^−1^ min^−1^ to 900 μmol mol^−1^. SS curves were collected starting at a reference CO_2_ concentration of 380 μmol mol^−1^, then proceeding through the following progression conditions 285, 190, 145, 100, 50, 380, 475, 570, 665, 760, 960,1160, 1460, 1760, 1960 μmol mol^−1^ CO_2_.

### Analysis

RACiR data were calibrated using an empty leaf chamber run taken within 1 hr using the R package {racir} (Stinziano et al. 2019a; Stinziano, 2020). *A/C*_i_, curves were fit using the fitaci() command from the {plantecophys} package (Duursma, 2015) in R statistical software (R Core Team, 2019, version 3.6.2), with options selected to fit triose phosphate utilization (*TPU*) limitation rate and with no temperature correction. We extracted maximum rubisco carboxylation capacity (*V*_cmax_), maximum electron transport rate under saturating light (*J*_1200_), *TPU*, fitted respiration (*R*_d_), the CO_2_ to RuBP transition *C*_i_ (*C*_itrans1_), and the RuBP to TPU transition *C*_i_ (*C*_itrans2_) for analysis. To assess a success rate, curves where more than half the data points had *g*_sw_ <0.08 μmol m^−2^ s^−1^ were considered to have failed.

### Statistics

The resulting parameters from the fit curves were subjected to paired Student’s *t*-tests to test for differences between steady state and RACiR curves after all points below the threshold had been removed.

### Data and Code

Data and code are available in supplementary files S1, S2, and S3.

## Results

### Success rate of A/C_i_ curves

No RACiR curves failed when beginning at *g*_sw_ >0.08 μmol m^−2^ s^−1^, however three SS curves out of 19 failed. Two of these curves lacked RuBP limitations (i.e. fitted *J*_1200_ = 1,000,000 - see source code of Duursma, 2015 for an explanation) or generated unreasonable estimates of *C*_itrans1_ (2,000 μmol mol^−1^) (Figure 1).

**Figure 1.**
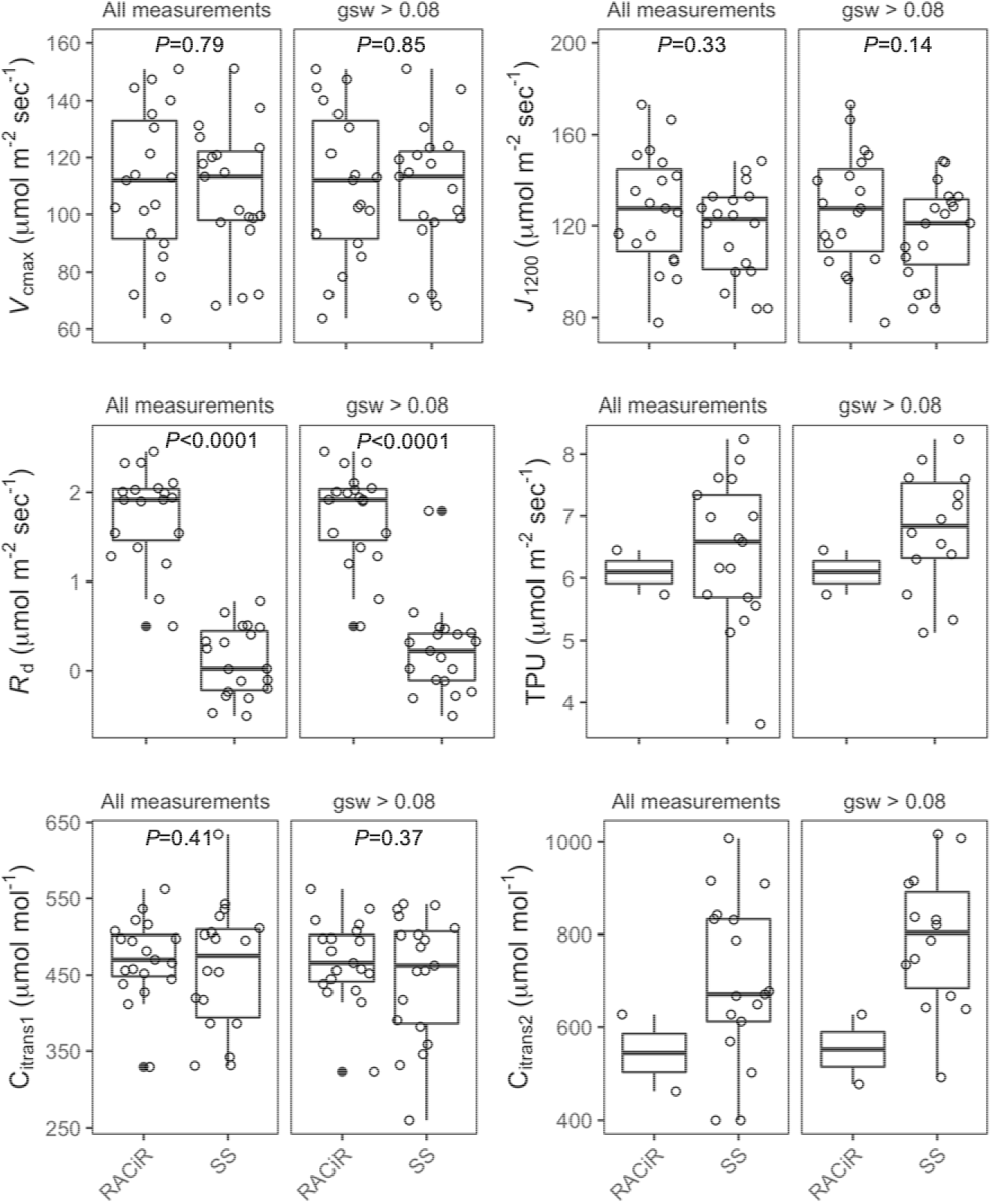
Box plots of *V*_cmax_, *J*_1200_, fitted *R*_d_, *TPU*, and transitions points of limitation states of steady state (SS) and RACiR *A/C*_i_ curves of *Citrus x sinensis* leaves with all data included or points with *g* < 0.08 μmol m^−2^ s^−1^ removed. *C*_i_ is the intercellular concentration of CO_2_. *C*_itrans1_ represents the transition between CO_2_ and RuBP limitations (i.e. *V*_max_ and *J*_1200_). *C*_itrans2_ is the transition from RuBP to TPU limitations (i.e. *J*_1200_ and *TPU*). To enable visualization one point measuring 2000 μmol mol^−1^ was removed from the SS All measurements group. Similarly, one *J*_1200_ estimate of 1,000,000 μmol m^−2^ s^−1^ was removed from the SS All measurements group (this value acts as a placeholder for the analysis and indicates no RuBP limitation, see Duursma, 2015). Open circles indicate individual data points. Closed circles represent distributions of outliers as determined by 95% confidence intervals. P-values indicate the probability that the SS and RACiR are not the same using a paired t-test assessing only the groups with *g*_sw_ above the threshold. All t-tests had 18 degrees of freedom.

### RACiR versus steady-state A/C_i_ curves

Overall, results from SS and RACiR curves were similar (Fig. 1). Notable differences are the lower *R*_d_ estimates by SS than by RACiR, and the small number of *TPU* estimates produced by RACiR, not surprising considering the preset range of leaf surface CO_2_ concentration (*C*_a_) values for RACiR.

### Stomatal limitations on gas exchange

Considering *A/C*_i_ curves (Fig. 2), three curves reduced *A* noticeably as *C*_i_ exceeded 700 μmol mol^−1^. However, a larger number of curves show *g*_sw_ dropping below our threshold of 0.08 μmol m^−2^ s^1^ as *C*_i_ increases *or* decreases relative to the “ambient” range of 380 μmol mol^−1^. This stomatal sensitivity to changes in *C*_a_ has implications for the validity of the resulting estimates based on curve fitting. Most notably, when points with *g*_sw_ <0.08 μmol m^−2^ s^−1^ were removed, estimates of *TPU* and the *C*_i_ of the RuBP-TPU limitation transition shifted upward for the SS curves. Additionally, variation between individual leaves in the responses to *C*_a_, resulted in a wide range of maximum *C*_i_ values in SS curves, and the minimum *g*_sw_ values of most curves were below 0.08 μmol m^−2^ s^−1^ (Fig. 2).

**Figure 2.**
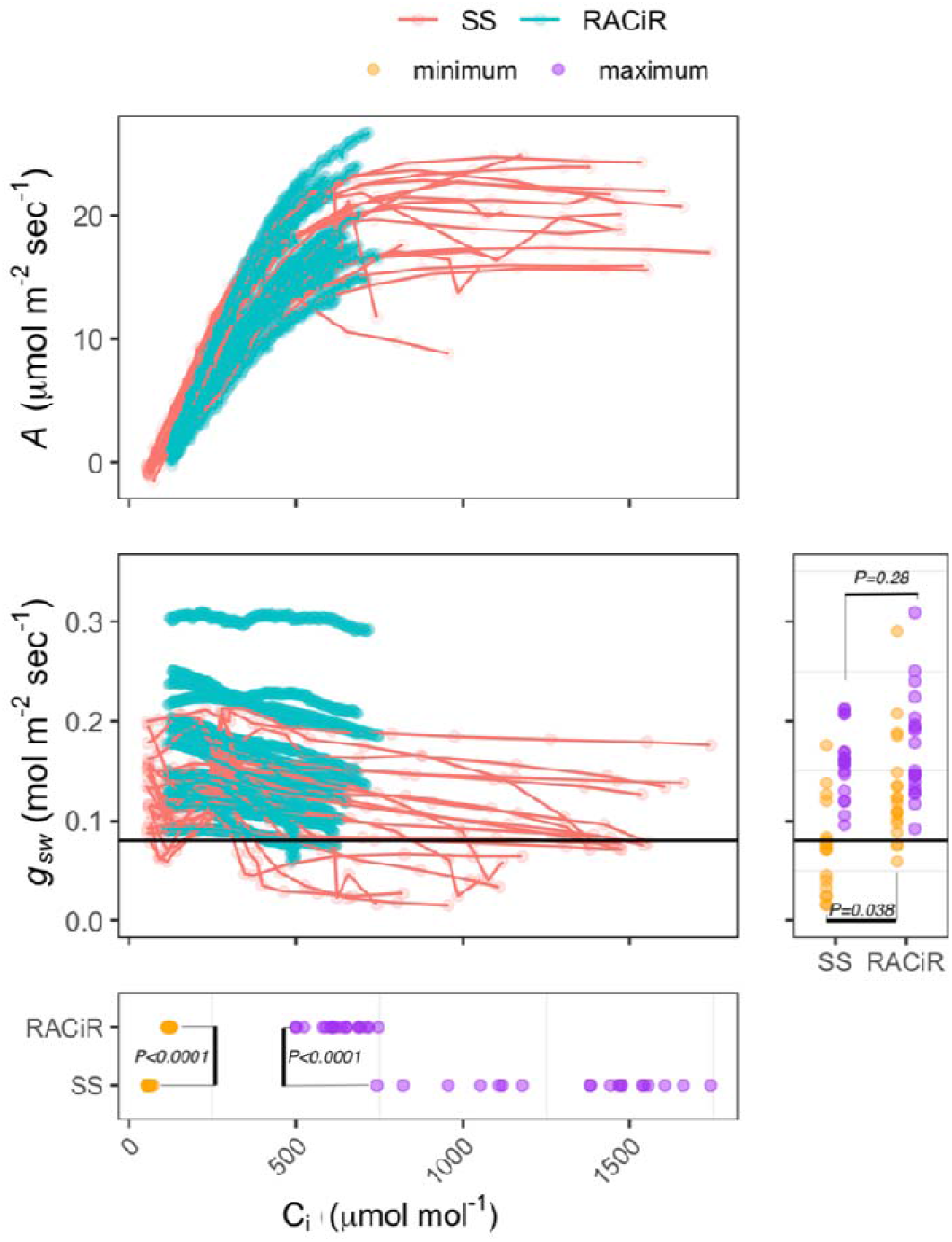
Plots of net CO_2_ assimilation (*A*) and stomatal conductance (*g*_sw_) by intercellular CO_2_ concentration (*C*_i_), and distributions of maximum and minimum *g*_sw_ and *C*_i_ of leaves of *Citrus x sinensis* using steady state (SS) or RACiR methods. Black horizontal line represents the *g*_sw_ threshold of 0.08 μmol m^−2^ s^−1^. *P*-values indicate results of a t-test with 18 degrees of freedom.

## Discussion

Overall, results of *A/C*_i_ curve fits measured by steady-state or RACiR methods are similar with more precise estimates of *C*_itrans1_ for RACiR (Fig. 1), confirming previous studies that the fitted parameters *V*_cmax_ and *J*_max_ are statistically equivalent (Stinziano et al., 2017; Coursolle et al., 2019; Lawrence et al., 2019; Stinziano et al., 2019b). The single difference was found in the estimation of *R*_d_, where the mean was approximately 1.75 ± 0.52 μmol m^−2^ s^−1^ for RACiR and 0.22 ± 0.5 μmol m^−2^ s^−1^ for the steady-state curves. On the same variety in a different study we found direct measurement of *R*_d_ to be 2.35 ± 0.11 μmol m^−2^ s^−1^ (Vincent et al., *unpublished data*).

Despite the similar resulting parameters, steady state curves are measured slowly relative to the time scale of leaf CO_2_ attenuation of *g*_sw_ (Merilo et al., 2015). This leads to the reduction of *g*_sw_ in some curves below the threshold required for reliable gas exchange measurements. This can particularly affect the estimation of *TPU*, as stomatal attenuation may induce changes in *A* that are limited by stomatal conductance, rather than by enzymatic limitations despite the similarity in shape of the curve to the expected shape with TPU limitations. We observed g_sw_ falling below a critical threshold in 3 of 19 cases for the steady-state curves compared to 0 cases for the RACiR curves. The likely explanation for these differences is due to the higher minimum and maximum *g*_sw_ in the RACiR curves versus the steady-state curves, creating a larger buffer and less time for *g*_sw_ to fall before reaching the critical threshold. We do note, however, that the traditional approach to *A/C*_i_ curves (starting near atmospheric CO_2_, going down, then up; e.g. Busch, 2018) may not be the best option for maximizing *g*_sw_ during the steady state, as starting at low CO_2_ and monotonically increasing CO_2_ may result in better *A/C*_i_ estimates (Sharkey, 2019), potentially avoiding the *g*_sw_ crash that can occur in *Citrus*.

To assess the *A/C*_i_ relationship, it is ideal to achieve a wide range of *C*_i_ values. The RACiR methods presented here did not achieve as wide a range of *C*_i_ as the SS methods. However, the very broad range of maximum *C*_i_ values demonstrates that the control of SS methods over this range is limited in such low conductance situations (Fig. 2). Additionally, depending on the species-specific responses, RACiR methods could continue to higher *C*_i_ values than those presented here, as the majority of curves ended with minimum *g*_sw_ well above the minimum threshold. In this case, the advantage of RACiR lies in that its short time “beats” the speed of stomatal attenuation, affording the ability to avoid stomatal impacts on the shape of the *A/C*_i_ curve. Additionally, we speculate that the speed of RACiR curves may allow for better estimation of *TPU* at high CO_2_, as such limitations are known to be transient and cryptic (Sharkey, 2019).

Therefore, based on our data, we recommend the use of RACiR in situations and species where rapid stomatal closure can limit the application of steady-state *A/C*_i_ curves.

## Supporting information

Supplement 3 results

