## Supplementary material for "Racing against stomatal attenuation: rapid CO_2_ response curves more reliably estimate photosynthetic capacity than steady state curves in a low conductance species": S1 code

#Loading packages to get started

#acifit <- fitaci(data, varnames = list(ALEAF = "A", Tleaf = "TleafCnd", Ci = "Ci", PPFD = "Qin"), fitmethod = "bilinear", Tcorrect = FALSE, fitTPU = TRUE)

library(processx)

library(devtools)

library(ggplotgui)

library(lubridate)

library(plantecophys)

library(Rcpp)

library(rmarkdown)

library(tidyverse)

library(racir)

library(reshape2)

library(patchwork)

library(ggplot2)

setwd("/Directory/subdirectory")

\#####

#RACiR curves

\#####

\######

#Tree 9

\#####

data<-read_6800("2020-03-19-RACiRtree9leaf1", skiplines=61)

#Check calibraton

racircalcheck(calfile="2020-03-19-RACiRtree5empty", skiplines = 61)

#nonlinearities on first check, remove values above 150

racircalcheck(calfile="2020-03-19-RACiRtree5empty", skiplines = 61, mincut=150)

#Calibrate file

racircal(calfile = "2020-03-19-RACiRtree5empty", datafile = "2020-03-19-RACiRtree9leaf1", skiplines = 61, mincut = 150)

data.R.9 <- read.csv("2020-03-19-RACiRtree9leaf1.csv",header = TRUE)

data.R.9$Type="RACiR"

data.R.9$Tree=9

acifit <- fitaci(data.R.9, varnames = list(ALEAF = "Acor", Tleaf = "TleafCnd",

Ci = "Cicor",

PPFD = "Qin"),

fitmethod = "bilinear" ,

Tcorrect = FALSE,

fitTPU = TRUE)

acifit$Ci_transition

acifit$Ci_transition2

plot(acifit)

coef(acifit)

\######

#Tree 11

\#####

data<-read_6800("2020-03-19-RACiRtree11leaf1", skiplines=61)

#Check calibraton

racircalcheck(calfile="2020-03-19-RACiRtree5empty", skiplines = 61)

#nonlinearities on first check, remove values above 150

racircalcheck(calfile="2020-03-19-RACiRtree5empty", skiplines = 61, mincut=150)

#Calibrate file

racircal(calfile = "2020-03-19-RACiRtree5empty", datafile = "2020-03-19-RACiRtree11leaf1", skiplines = 61, mincut = 150)

data.R.11 <- read.csv("2020-03-19-RACiRtree11leaf1.csv",header = TRUE)

data.R.11$Type="RACiR"

data.R.11$Tree=11

acifit <- fitaci(data.R.11, varnames = list(ALEAF = "Acor", Tleaf = "TleafCnd",

Ci = "Cicor",

PPFD = "Qin"),

fitmethod = "bilinear" ,

Tcorrect = FALSE,

fitTPU = TRUE)

acifit$Ci_transition

acifit$Ci_transition2

plot(acifit)

coef(acifit)

\######

#Tree 12

\#####

data<-read_6800("2020-03-19-RACiRtree12leaf1", skiplines=61)

#Check calibraton

racircalcheck(calfile="2020-03-19-RACiRtree5empty", skiplines = 61)

#nonlinearities on first check, remove values above 150

racircalcheck(calfile="2020-03-19-RACiRtree5empty", skiplines = 61, mincut=150)

#Calibrate file

racircal(calfile = "2020-03-19-RACiRtree5empty", datafile = "2020-03-19-RACiRtree12leaf1", skiplines = 61, mincut = 150)

data.R.12 <- read.csv("2020-03-19-RACiRtree12leaf1.csv",header = TRUE)

data.R.12$Type="RACiR"

data.R.12$Tree=12

acifit <- fitaci(data.R.12, varnames = list(ALEAF = "Acor", Tleaf = "TleafCnd",

Ci = "Cicor",

PPFD = "Qin"),

fitmethod = "bilinear" ,

Tcorrect = FALSE,

fitTPU = TRUE)

acifit$Ci_transition

acifit$Ci_transition2

plot(acifit)

coef(acifit)

\######

#Tree 13

\#####

data<-read_6800("2020-03-19-RACiRtree13leaf1", skiplines=61)

#Check calibraton

racircalcheck(calfile="2020-03-19-RACiRtree5empty", skiplines = 61)

#nonlinearities on first check, remove values above 150

racircalcheck(calfile="2020-03-19-RACiRtree5empty", skiplines = 61, mincut=150)

#Calibrate file

racircal(calfile = "2020-03-19-RACiRtree5empty", datafile = "2020-03-19-RACiRtree13leaf1", skiplines = 61, mincut = 150)

data.R.13 <- read.csv("2020-03-19-RACiRtree13leaf1.csv",header = TRUE)

data.R.13$Type="RACiR"

data.R.13$Tree=13

acifit <- fitaci(data.R.13, varnames = list(ALEAF = "Acor", Tleaf = "TleafCnd",

Ci = "Cicor",

PPFD = "Qin"),

fitmethod = "bilinear" ,

Tcorrect = FALSE,

fitTPU = TRUE)

acifit$Ci_transition

acifit$Ci_transition2

plot(acifit)

coef(acifit)

\######

#Tree 14

\#####

data<-read_6800("2020-03-19-RACiRtree14leaf1", skiplines=61)

#Check calibraton

racircalcheck(calfile="2020-03-19-RACiRtree5empty", skiplines = 61)

#nonlinearities on first check, remove values above 150

racircalcheck(calfile="2020-03-19-RACiRtree5empty", skiplines = 61, mincut=150)

#Calibrate file

racircal(calfile = "2020-03-19-RACiRtree5empty", datafile = "2020-03-19-RACiRtree14leaf1", skiplines = 61, mincut = 150)

data.R.14 <- read.csv("2020-03-19-RACiRtree14leaf1.csv",header = TRUE)

data.R.14$Type="RACiR"

data.R.14$Tree=14

acifit <- fitaci(data.R.14, varnames = list(ALEAF = "Acor", Tleaf = "TleafCnd",

Ci = "Cicor",

PPFD = "Qin"),

fitmethod = "bilinear" ,

Tcorrect = FALSE,

fitTPU = TRUE)

acifit$Ci_transition

acifit$Ci_transition2

plot(acifit)

coef(acifit)

\######

#Tree 15

\#####

data<-read_6800("2020-03-20-RACiRtree15leaf1", skiplines=61)

#Check calibraton

racircalcheck(calfile="2020-03-20-RACiRleaf6empty", skiplines = 61)

#nonlinearities on first check, remove values above 150

racircalcheck(calfile="2020-03-20-RACiRleaf6empty", skiplines = 61, mincut=150)

#Calibrate file

racircal(calfile = "2020-03-20-RACiRleaf6empty", datafile = "2020-03-20-RACiRtree15leaf1", skiplines = 61, mincut = 150)

data.R.15 <- read.csv("2020-03-20-RACiRtree15leaf1.csv",header = TRUE)

data.R.15$Type="RACiR"

data.R.15$Tree=15

acifit <- fitaci(data.R.15, varnames = list(ALEAF = "Acor", Tleaf = "TleafCnd",

Ci = "Cicor",

PPFD = "Qin"),

fitmethod = "bilinear" ,

Tcorrect = FALSE,

fitTPU = TRUE)

acifit$Ci_transition

acifit$Ci_transition2

plot(acifit)

coef(acifit)

\######

#Tree 16

\#####

data<-read_6800("2020-03-20-RACiRtree16leaf1", skiplines=61)

#Check calibraton

racircalcheck(calfile="2020-03-20-RACiRleaf6empty", skiplines = 61)

#nonlinearities on first check, remove values above 150

racircalcheck(calfile="2020-03-20-RACiRleaf6empty", skiplines = 61, mincut=150)

#Calibrate file

racircal(calfile = "2020-03-20-RACiRleaf6empty", datafile = "2020-03-20-RACiRtree16leaf1", skiplines = 61, mincut = 150)

data.R.16 <- read.csv("2020-03-20-RACiRtree16leaf1.csv",header = TRUE)

data.R.16$Type="RACiR"

data.R.16$Tree=16

acifit <- fitaci(data.R.16, varnames = list(ALEAF = "Acor", Tleaf = "TleafCnd",

Ci = "Cicor",

PPFD = "Qin"),

fitmethod = "bilinear" ,

Tcorrect = FALSE,

fitTPU = TRUE)

acifit$Ci_transition

acifit$Ci_transition2

plot(acifit)

coef(acifit)

\######

#Tree 17

\#####

data<-read_6800("2020-03-23-RACiRtree17leaf1", skiplines=61)

#Check calibraton

racircalcheck(calfile="2020-03-23-RACiRtree17empty", skiplines = 61)

#nonlinearities on first check, remove values above 150

racircalcheck(calfile="2020-03-23-RACiRtree17empty", skiplines = 61, mincut=150)

#Calibrate file

racircal(calfile = "2020-03-23-RACiRtree17empty", datafile = "2020-03-23-RACiRtree17leaf1", skiplines = 61, mincut = 150)

data.R.17 <- read.csv("2020-03-23-RACiRtree17leaf1.csv",header = TRUE)

data.R.17$Type="RACiR"

data.R.17$Tree=17

acifit <- fitaci(data.R.17, varnames = list(ALEAF = "Acor", Tleaf = "TleafCnd",

Ci = "Cicor",

PPFD = "Qin"),

fitmethod = "bilinear" ,

Tcorrect = FALSE,

fitTPU = TRUE)

acifit$Ci_transition

acifit$Ci_transition2

plot(acifit)

coef(acifit)

\######

#Tree 18

\#####

data<-read_6800("2020-03-23-RACiRtree18leaf1", skiplines=61)

#Check calibraton

racircalcheck(calfile="2020-03-23-RACiRtree17empty", skiplines = 61)

#nonlinearities on first check, remove values above 150

racircalcheck(calfile="2020-03-23-RACiRtree17empty", skiplines = 61, mincut=150)

#Calibrate file

racircal(calfile = "2020-03-23-RACiRtree17empty", datafile = "2020-03-23-RACiRtree18leaf1", skiplines = 61, mincut = 150)

data.R.18 <- read.csv("2020-03-23-RACiRtree18leaf1.csv",header = TRUE)

data.R.18$Type="RACiR"

data.R.18$Tree=18

acifit <- fitaci(data.R.18, varnames = list(ALEAF = "Acor", Tleaf = "TleafCnd",

Ci = "Cicor",

PPFD = "Qin"),

fitmethod = "bilinear" ,

Tcorrect = FALSE,

fitTPU = TRUE)

acifit$Ci_transition

acifit$Ci_transition2

plot(acifit)

coef(acifit)

\######

#Tree 19

\#####

plot(acifit)

coef(acifit)

\######

#Tree 20

\#####

data<-read_6800("2020-03-23-RACiRtree20leaf1", skiplines=61)

#Check calibraton

racircalcheck(calfile="2020-03-23-RACiRtree17empty", skiplines = 61)

#nonlinearities on first check, remove values above 150

racircalcheck(calfile="2020-03-23-RACiRtree17empty", skiplines = 61, mincut=150)

#Calibrate file

racircal(calfile = "2020-03-23-RACiRtree17empty", datafile = "2020-03-23-RACiRtree20leaf1", skiplines = 61, mincut = 150)

data.R.20 <- read.csv("2020-03-23-RACiRtree20leaf1.csv",header = TRUE)

data.R.20$Type="RACiR"

data.R.20$Tree=20

acifit <- fitaci(data.R.20, varnames = list(ALEAF = "Acor", Tleaf = "TleafCnd",

Ci = "Cicor",

PPFD = "Qin"),

fitmethod = "bilinear" ,

Tcorrect = FALSE,

fitTPU = TRUE)

acifit$Ci_transition

acifit$Ci_transition2

plot(acifit)

coef(acifit)

\######

#Tree 21

\#####

data<-read_6800("2020-04-09-RACiRtree21leaf1", skiplines=61)

#Check calibraton

racircalcheck(calfile="2020-04-09-RACiRtree20empty", skiplines = 61)

#nonlinearities on first check, remove values above 150

racircalcheck(calfile="2020-04-09-RACiRtree20empty", skiplines = 61, mincut=150)

#Calibrate file

racircal(calfile = "2020-04-09-RACiRtree20empty", datafile = "2020-04-09-RACiRtree21leaf1", skiplines = 61, mincut = 150)

data.R.21 <- read.csv("2020-04-09-RACiRtree21leaf1.csv",header = TRUE)

data.R.21$Type="RACiR"

data.R.21$Tree=21

acifit <- fitaci(data.R.21, varnames = list(ALEAF = "Acor", Tleaf = "TleafCnd",

Ci = "Cicor",

PPFD = "Qin"),

fitmethod = "bilinear" ,

Tcorrect = FALSE,

fitTPU = TRUE)

acifit$Ci_transition

acifit$Ci_transition2

plot(acifit)

coef(acifit)

\######

#Tree 22

\#####

data<-read_6800("2020-04-09-RACiRtree22leaf1", skiplines=61)

#Check calibraton

racircalcheck(calfile="2020-04-09-RACiRtree20empty", skiplines = 61)

#nonlinearities on first check, remove values above 150

racircalcheck(calfile="2020-04-09-RACiRtree20empty", skiplines = 61, mincut=150)

#Calibrate file

racircal(calfile = "2020-04-09-RACiRtree20empty", datafile = "2020-04-09-RACiRtree22leaf1", skiplines = 61, mincut = 150)

data.R.22 <- read.csv("2020-04-09-RACiRtree22leaf1.csv",header = TRUE)

data.R.22$Type="RACiR"

data.R.22$Tree=22

acifit <- fitaci(data.R.22, varnames = list(ALEAF = "Acor", Tleaf = "TleafCnd",

Ci = "Cicor",

PPFD = "Qin"),

fitmethod = "bilinear" ,

Tcorrect = FALSE,

fitTPU = TRUE)

acifit$Ci_transition

acifit$Ci_transition2

plot(acifit)

coef(acifit)

\######

#Tree 23

\#####

data<-read_6800("2020-04-09-RACiRtree23leaf1", skiplines=61)

#Check calibraton

racircalcheck(calfile="2020-04-09-RACiRtree20empty", skiplines = 61)

#nonlinearities on first check, remove values above 150

racircalcheck(calfile="2020-04-09-RACiRtree20empty", skiplines = 61, mincut=150)

#Calibrate file

racircal(calfile = "2020-04-09-RACiRtree20empty", datafile = "2020-04-09-RACiRtree23leaf1", skiplines = 61, mincut = 150)

data.R.23 <- read.csv("2020-04-09-RACiRtree23leaf1.csv",header = TRUE)

data.R.23$Type="RACiR"

data.R.23$Tree=23

acifit <- fitaci(data.R.23, varnames = list(ALEAF = "Acor", Tleaf = "TleafCnd",

Ci = "Cicor",

PPFD = "Qin"),

fitmethod = "bilinear" ,

Tcorrect = FALSE,

fitTPU = TRUE)

acifit$Ci_transition

acifit$Ci_transition2

plot(acifit)

coef(acifit)

\######

#Tree 24

\#####

data<-read_6800("2020-04-09-RACiRtree24leaf1", skiplines=61)

#Check calibraton

racircalcheck(calfile="2020-04-09-RACiRtree20empty", skiplines = 61)

#nonlinearities on first check, remove values above 150

racircalcheck(calfile="2020-04-09-RACiRtree20empty", skiplines = 61, mincut=150)

#Calibrate file

racircal(calfile = "2020-04-09-RACiRtree20empty", datafile = "2020-04-09-RACiRtree24leaf1", skiplines = 61, mincut = 150)

data.R.24 <- read.csv("2020-04-09-RACiRtree24leaf1.csv",header = TRUE)

data.R.24$Type="RACiR"

data.R.24$Tree=24

acifit <- fitaci(data.R.24, varnames = list(ALEAF = "Acor", Tleaf = "TleafCnd",

Ci = "Cicor",

PPFD = "Qin"),

fitmethod = "bilinear" ,

Tcorrect = FALSE,

fitTPU = TRUE)

acifit$Ci_transition

acifit$Ci_transition2

plot(acifit)

coef(acifit)

\######

#Tree 2

\#####

data<-read_6800("2020-06-16-RACiRtree2leaf1", skiplines=61)

#Check calibraton

racircalcheck(calfile="2020-06-16-RACiRleaf2emty", skiplines = 61)

#nonlinearities on first check, remove values above 150

racircalcheck(calfile="2020-06-16-RACiRleaf2emty", skiplines = 61, mincut=150)

#Calibrate file

racircal(calfile = "2020-06-16-RACiRleaf2emty", datafile = "2020-06-16-RACiRtree2leaf1", skiplines = 61, mincut = 150)

data.R.2 <- read.csv("2020-06-16-RACiRtree2leaf1.csv",header = TRUE)

data.R.2$Type="RACiR"

data.R.2$Tree=2

acifit <- fitaci(data.R.2, varnames = list(ALEAF = "Acor", Tleaf = "TleafCnd",

Ci = "Cicor",

PPFD = "Qin"),

fitmethod = "bilinear" ,

Tcorrect = FALSE,

fitTPU = TRUE)

acifit$Ci_transition

acifit$Ci_transition2

plot(acifit)

coef(acifit)

\######

#Tree 3

\#####

data<-read_6800("2020-06-16-RACiRtree3leaf1", skiplines=61)

#Check calibraton

racircalcheck(calfile="2020-06-16-RACiRleaf2emty", skiplines = 61)

#nonlinearities on first check, remove values above 150

racircalcheck(calfile="2020-06-16-RACiRleaf2emty", skiplines = 61, mincut=150)

#Calibrate file

racircal(calfile = "2020-06-16-RACiRleaf2emty", datafile = "2020-06-16-RACiRtree3leaf1", skiplines = 61, mincut = 150)

data.R.3 <- read.csv("2020-06-16-RACiRtree3leaf1.csv",header = TRUE)

data.R.3$Type="RACiR"

data.R.3$Tree=3

acifit <- fitaci(data.R.3, varnames = list(ALEAF = "Acor", Tleaf = "TleafCnd",

Ci = "Cicor",

PPFD = "Qin"),

fitmethod = "bilinear" ,

Tcorrect = FALSE,

fitTPU = TRUE)

acifit$Ci_transition

acifit$Ci_transition2

plot(acifit)

coef(acifit)

\######

#Tree 4

\#####

data<-read_6800("2020-06-16-RACiRtree4leaf1", skiplines=61)

#Check calibraton

racircalcheck(calfile="2020-06-16-RACiRleaf2emty", skiplines = 61)

#nonlinearities on first check, remove values above 150

racircalcheck(calfile="2020-06-16-RACiRleaf2emty", skiplines = 61, mincut=150)

#Calibrate file

racircal(calfile = "2020-06-16-RACiRleaf2emty", datafile = "2020-06-16-RACiRtree4leaf1", skiplines = 61, mincut = 150)

data.R.4 <- read.csv("2020-06-16-RACiRtree4leaf1.csv",header = TRUE)

data.R.4$Type="RACiR"

data.R.4$Tree=4

acifit <- fitaci(data.R.4, varnames = list(ALEAF = "Acor", Tleaf = "TleafCnd",

Ci = "Cicor",

PPFD = "Qin"),

fitmethod = "bilinear" ,

Tcorrect = FALSE,

fitTPU = TRUE)

acifit$Ci_transition

acifit$Ci_transition2

plot(acifit)

coef(acifit)

\######

#Tree 5

\#####

data<-read_6800("2020-06-16-RACiRtree5leaf1", skiplines=61)

#Check calibraton

racircalcheck(calfile="2020-06-16-RACiRleaf2emty", skiplines = 61)

#nonlinearities on first check, remove values above 150

racircalcheck(calfile="2020-06-16-RACiRleaf2emty", skiplines = 61, mincut=150)

#Calibrate file

racircal(calfile = "2020-06-16-RACiRleaf2emty", datafile = "2020-06-16-RACiRtree5leaf1", skiplines = 61, mincut = 150)

data.R.5 <- read.csv("2020-06-16-RACiRtree5leaf1.csv",header = TRUE)

data.R.5$Type="RACiR"

data.R.5$Tree="5"

acifit <- fitaci(data.R.5, varnames = list(ALEAF = "Acor", Tleaf = "TleafCnd",

Ci = "Cicor",

PPFD = "Qin"),

fitmethod = "bilinear" ,

Tcorrect = FALSE,

fitTPU = TRUE)

acifit$Ci_transition

acifit$Ci_transition2

plot(acifit)

coef(acifit)

\#####

#ACi steady state curves

\#####

\######

#Tree 4

\#####

data.A.4<-read_6800("2019-10-14-ACiRtree4leef1", skiplines=53)

data.A.4$Type="Aci"

data.A.4$Tree=4

acifit <- fitaci(data.A.4, varnames = list(ALEAF = "A", Tleaf = "TleafCnd",

Ci = "Ci",

PPFD = "Qin"),

fitmethod = "bilinear" ,

Tcorrect = FALSE,

fitTPU = TRUE)

plot(acifit)

coef(acifit)

\######

#Tree 9

\#####

data.A.9<-read_6800("2020-03-19-ACitree9leaf1", skiplines=61)

data.A.9$Type="Aci"

data.A.9$Tree=9

acifit <- fitaci(data.A.9, varnames = list(ALEAF = "A", Tleaf = "TleafCnd",

Ci = "Ci",

PPFD = "Qin"),

fitmethod = "bilinear" ,

Tcorrect = FALSE,

fitTPU = TRUE)

plot(acifit)

coef(acifit)

acifit$Ci_transition

acifit$Ci_transition2

\######

#Tree 11

\#####

data.A.11<-read_6800("2020-03-19-ACitree11leaf1", skiplines=61)

data.A.11$Type="Aci"

data.A.11$Tree=11

acifit <- fitaci(data.A.11, varnames = list(ALEAF = "A", Tleaf = "TleafCnd",

Ci = "Ci",

PPFD = "Qin"),

fitmethod = "bilinear" ,

Tcorrect = FALSE,

fitTPU = TRUE)

plot(acifit)

coef(acifit)

\######

#Tree 12

\#####

data.A.12<-read_6800("2020-03-19-ACitree12leaf1", skiplines=61)

data.A.12$Type="Aci"

data.A.12$Tree=12

acifit <- fitaci(data.A.12, varnames = list(ALEAF = "A", Tleaf = "TleafCnd",

Ci = "Ci",

PPFD = "Qin"),

fitmethod = "bilinear" ,

Tcorrect = FALSE,

fitTPU = TRUE)

plot(acifit)

coef(acifit)

acifit$Ci_transition

acifit$Ci_transition2

\######

#Tree 13

\#####

data.A.13<-read_6800("2020-03-19-ACitree13leaf1", skiplines=61)

data.A.13$Type="Aci"

data.A.13$Tree=13

acifit <- fitaci(data.A.13, varnames = list(ALEAF = "A", Tleaf = "TleafCnd",

Ci = "Ci",

PPFD = "Qin"),

fitmethod = "bilinear" ,

Tcorrect = FALSE,

fitTPU = TRUE)

plot(acifit)

coef(acifit)

acifit$Ci_transition

acifit$Ci_transition2

\######

#Tree 14

\#####

data.A.14<-read_6800("2020-03-19-ACitree14leaf1", skiplines=61)

data.A.14$Type="Aci"

data.A.14$Tree=14

acifit <- fitaci(data.A.14, varnames = list(ALEAF = "A", Tleaf = "TleafCnd",

Ci = "Ci",

PPFD = "Qin"),

fitmethod = "bilinear" ,

Tcorrect = FALSE,

fitTPU = TRUE)

plot(acifit)

coef(acifit)

acifit$Ci_transition

acifit$Ci_transition2

\######

#Tree 15

\#####

data.A.15<-read_6800("2020-03-20-ACitree15leaf1", skiplines=61)

data.A.15$Type="Aci"

data.A.15$Tree=15

acifit <- fitaci(data.A.15, varnames = list(ALEAF = "A", Tleaf = "TleafCnd",

Ci = "Ci",

PPFD = "Qin"),

fitmethod = "bilinear" ,

Tcorrect = FALSE,

fitTPU = TRUE)

plot(acifit)

coef(acifit)

acifit$Ci_transition

acifit$Ci_transition2

\######

#Tree 16

\#####

data.A.16<-read_6800("2020-03-20-ACitree16leaf1", skiplines=61)

data.A.16$Type="Aci"

data.A.16$Tree=16

acifit <- fitaci(data.A.16, varnames = list(ALEAF = "A", Tleaf = "TleafCnd",

Ci = "Ci",

PPFD = "Qin"),

fitmethod = "bilinear" ,

Tcorrect = FALSE,

fitTPU = TRUE)

plot(acifit)

coef(acifit)

acifit$Ci_transition

acifit$Ci_transition2

\######

#Tree 17

\#####

data.A.17<-read_6800("2020-03-23-ACitree17leaf1", skiplines=61)

data.A.17$Type="Aci"

data.A.17$Tree=17

acifit <- fitaci(data.A.17, varnames = list(ALEAF = "A", Tleaf = "TleafCnd",

Ci = "Ci",

PPFD = "Qin"),

fitmethod = "bilinear" ,

Tcorrect = FALSE,

fitTPU = TRUE)

plot(acifit)

coef(acifit)

acifit$Ci_transition

acifit$Ci_transition2

\######

#Tree 18

\#####

data.A.18<-read_6800("2020-03-23-ACitree18leaf1", skiplines=61)

data.A.18$Type="Aci"

data.A.18$Tree=18

acifit <- fitaci(data.A.18, varnames = list(ALEAF = "A", Tleaf = "TleafCnd",

Ci = "Ci",

PPFD = "Qin"),

fitmethod = "bilinear" ,

Tcorrect = FALSE,

fitTPU = TRUE)

plot(acifit)

coef(acifit)

acifit$Ci_transition

acifit$Ci_transition2

\######

#Tree 19

\#####

data.A.19<-read_6800("2020-03-23-ACitree19leaf1", skiplines=61)

data.A.19$Type="Aci"

data.A.19$Tree=19

acifit <- fitaci(data.A.19, varnames = list(ALEAF = "A", Tleaf = "TleafCnd",

Ci = "Ci",

PPFD = "Qin"),

fitmethod = "bilinear" ,

Tcorrect = FALSE,

fitTPU = TRUE)

plot(acifit)

coef(acifit)

acifit$Ci_transition

acifit$Ci_transition2

\######

#Tree 20

\#####

data.A.20<-read_6800("2020-03-23-ACitree20leaf1", skiplines=61)

data.A.20$Type="Aci"

data.A.20$Tree=20

acifit <- fitaci(data.A.20, varnames = list(ALEAF = "A", Tleaf = "TleafCnd",

Ci = "Ci",

PPFD = "Qin"),

fitmethod = "bilinear" ,

Tcorrect = FALSE,

fitTPU = TRUE)

plot(acifit)

coef(acifit)

acifit$Ci_transition

acifit$Ci_transition2

\######

#Tree 21

\#####

data.A.21<-read_6800("2020-04-09-ACitree21leaf1", skiplines=61)

data.A.21$Type="Aci"

data.A.21$Tree=21

acifit <- fitaci(data.A.21, varnames = list(ALEAF = "A", Tleaf = "TleafCnd",

Ci = "Ci",

PPFD = "Qin"),

fitmethod = "bilinear" ,

Tcorrect = FALSE,

fitTPU = TRUE)

plot(acifit)

coef(acifit)

acifit$Ci_transition

acifit$Ci_transition2

\######

#Tree 22

\#####

data.A.22<-read_6800("2020-04-09-ACitree22leaf1", skiplines=61)

data.A.22$Type="Aci"

data.A.22$Tree=22

acifit <- fitaci(data.A.22, varnames = list(ALEAF = "A", Tleaf = "TleafCnd",

Ci = "Ci",

PPFD = "Qin"),

fitmethod = "bilinear" ,

Tcorrect = FALSE,

fitTPU = TRUE)

plot(acifit)

coef(acifit)

acifit$Ci_transition

acifit$Ci_transition2

\######

#Tree 23

\#####

data.A.23<-read_6800("2020-04-09-ACitree23leaf1", skiplines=61)

data.A.23$Type="Aci"

data.A.23$Tree=23

acifit <- fitaci(data.A.23, varnames = list(ALEAF = "A", Tleaf = "TleafCnd",

Ci = "Ci",

PPFD = "Qin"),

fitmethod = "bilinear" ,

Tcorrect = FALSE,

fitTPU = TRUE)

plot(acifit)

coef(acifit)

acifit$Ci_transition

acifit$Ci_transition2

\######

#Tree 24

\#####

data.A.24<-read_6800("2020-04-09-ACitree24leaf1", skiplines=61)

data.A.24$Type="Aci"

data.A.24$Tree=24

acifit <- fitaci(data.A.24, varnames = list(ALEAF = "A", Tleaf = "TleafCnd",

Ci = "Ci",

PPFD = "Qin"),

fitmethod = "bilinear" ,

Tcorrect = FALSE,

fitTPU = TRUE)

plot(acifit)

coef(acifit)

acifit$Ci_transition

acifit$Ci_transition2

\######

#Tree 2

\#####

data.A.2<-read_6800("2020-06-16-ACitree2leaf1", skiplines=61)

data.A.2$Type="Aci"

data.A.2$Tree=2

acifit <- fitaci(data.A.2, varnames = list(ALEAF = "A", Tleaf = "TleafCnd",

Ci = "Ci",

PPFD = "Qin"),

fitmethod = "bilinear" ,

Tcorrect = FALSE,

fitTPU = TRUE)

plot(acifit)

coef(acifit)

acifit$Ci_transition

acifit$Ci_transition2

\######

#Tree 3

\#####

data.A.3<-read_6800("2020-06-16-ACitree3leaf1", skiplines=61)

data.A.3$Type="Aci"

data.A.3$Tree=3

acifit <- fitaci(data.A.3, varnames = list(ALEAF = "A", Tleaf = "TleafCnd",

Ci = "Ci",

PPFD = "Qin"),

fitmethod = "bilinear" ,

Tcorrect = FALSE,

fitTPU = TRUE)

plot(acifit)

coef(acifit)

acifit$Ci_transition

acifit$Ci_transition2

\######

#Tree 4

\#####

data.A.4<-read_6800("2020-06-16-ACitree4leaf1", skiplines=61)

data.A.4$Type="Aci"

data.A.4$Tree=4

acifit <- fitaci(data.A.4, varnames = list(ALEAF = "A", Tleaf = "TleafCnd",

Ci = "Ci",

PPFD = "Qin"),

fitmethod = "bilinear" ,

Tcorrect = FALSE,

fitTPU = TRUE)

plot(acifit)

coef(acifit)

acifit$Ci_transition

acifit$Ci_transition2

\######

#Tree 5

\#####

data.A.5<-read_6800("2020-06-16-ACitree5leaf1", skiplines=61)

data.A.5$Type="Aci"

data.A.5$Tree=5

acifit <- fitaci(data.A.5, varnames = list(ALEAF = "A", Tleaf = "TleafCnd",

Ci = "Ci",

PPFD = "Qin"),

fitmethod = "bilinear" ,

Tcorrect = FALSE,

fitTPU = TRUE)

plot(acifit)

coef(acifit)

acifit$Ci_transition

acifit$Ci_transition2

\#####

#Combining all runs

\#####

A.2<-select(data.A.2, gsw,A,Ci,Type,Tree,TleafCnd,Qin,Ca)

A.3<-select(data.A.3, gsw,A,Ci,Type,Tree,TleafCnd,Qin,Ca)

A.4<-select(data.A.4, gsw,A,Ci,Type,Tree,TleafCnd,Qin,Ca)

A.5<-select(data.A.5, gsw,A,Ci,Type,Tree,TleafCnd,Qin,Ca)

A.9<-select(data.A.9, gsw,A,Ci,Type,Tree,TleafCnd,Qin,Ca)

A.11<-select(data.A.11, gsw,A,Ci,Type,Tree,TleafCnd,Qin,Ca)

A.12<-select(data.A.12, gsw,A,Ci,Type,Tree,TleafCnd,Qin,Ca)

A.13<-select(data.A.13, gsw,A,Ci,Type,Tree,TleafCnd,Qin,Ca)

A.14<-select(data.A.14, gsw,A,Ci,Type,Tree,TleafCnd,Qin,Ca)

A.15<-select(data.A.15, gsw,A,Ci,Type,Tree,TleafCnd,Qin,Ca)

A.16<-select(data.A.16, gsw,A,Ci,Type,Tree,TleafCnd,Qin,Ca)

A.17<-select(data.A.17, gsw,A,Ci,Type,Tree,TleafCnd,Qin,Ca)

A.18<-select(data.A.18, gsw,A,Ci,Type,Tree,TleafCnd,Qin,Ca)

A.19<-select(data.A.19, gsw,A,Ci,Type,Tree,TleafCnd,Qin,Ca)

A.20<-select(data.A.20, gsw,A,Ci,Type,Tree,TleafCnd,Qin,Ca)

A.21<-select(data.A.21, gsw,A,Ci,Type,Tree,TleafCnd,Qin,Ca)

A.22<-select(data.A.22, gsw,A,Ci,Type,Tree,TleafCnd,Qin,Ca)

A.23<-select(data.A.23, gsw,A,Ci,Type,Tree,TleafCnd,Qin,Ca)

A.24<-select(data.A.24, gsw,A,Ci,Type,Tree,TleafCnd,Qin,Ca)

R.2<-select(data.R.2, gsw,Acor,Cicor,Type,Tree,TleafCnd,Qin,Ca)

R.3<-select(data.R.3, gsw,Acor,Cicor,Type,Tree,TleafCnd,Qin,Ca)

R.4<-select(data.R.4, gsw,Acor,Cicor,Type,Tree,TleafCnd,Qin,Ca)

R.5<-select(data.R.5, gsw,Acor,Cicor,Type,Tree,TleafCnd,Qin,Ca)

R.9<-select(data.R.9, gsw,Acor,Cicor,Type,Tree,TleafCnd,Qin,Ca)

R.11<-select(data.R.11, gsw,Acor,Cicor,Type,Tree,TleafCnd,Qin,Ca)

R.12<-select(data.R.12, gsw,Acor,Cicor,Type,Tree,TleafCnd,Qin,Ca)

R.13<-select(data.R.13, gsw,Acor,Cicor,Type,Tree,TleafCnd,Qin,Ca)

R.14<-select(data.R.14, gsw,Acor,Cicor,Type,Tree,TleafCnd,Qin,Ca)

R.15<-select(data.R.15, gsw,Acor,Cicor,Type,Tree,TleafCnd,Qin,Ca)

R.16<-select(data.R.16, gsw,Acor,Cicor,Type,Tree,TleafCnd,Qin,Ca)

R.17<-select(data.R.17, gsw,Acor,Cicor,Type,Tree,TleafCnd,Qin,Ca)

R.18<-select(data.R.18, gsw,Acor,Cicor,Type,Tree,TleafCnd,Qin,Ca)

R.19<-select(data.R.19, gsw,Acor,Cicor,Type,Tree,TleafCnd,Qin,Ca)

R.20<-select(data.R.20, gsw,Acor,Cicor,Type,Tree,TleafCnd,Qin,Ca)

R.21<-select(data.R.21, gsw,Acor,Cicor,Type,Tree,TleafCnd,Qin,Ca)

R.22<-select(data.R.22, gsw,Acor,Cicor,Type,Tree,TleafCnd,Qin,Ca)

R.23<-select(data.R.23, gsw,Acor,Cicor,Type,Tree,TleafCnd,Qin,Ca)

R.24<-select(data.R.24, gsw,Acor,Cicor,Type,Tree,TleafCnd,Qin,Ca)

library(plyr)

A.df<-rbind(A.2,A.3,A.4,A.5,A.9,A.11,A.12,A.13,A.14,A.15,

A.16,A.17,A.18,A.19,A.20,A.21,A.22,

A.23,A.24)

R.df<-rbind(R.2,R.3,R.4,R.5,R.9,R.11,R.12,R.13,R.14,R.15,

R.16,R.17,R.18,R.19,R.20,R.21,R.22,

R.23,R.24)

R.df<-R.df %>% dplyr::rename(Ci=Cicor, A=Acor)

allcurves.df<-rbind(A.df,R.df)

allcurves.df<-read.table("/Directory/subdirectory/Total ACi data.csv", header = TRUE, sep = ",", dec=".", fill = TRUE)

head(allcurves.df)

\#####

#creating ID variables

\#####

allcurves.df$Measure<-allcurves.df$Type

allcurves.df$ID[allcurves.df$Tree=="2" & allcurves.df$Measure=="ACi"]<-"1"

allcurves.df$ID[allcurves.df$Tree=="3" & allcurves.df$Measure=="ACi"]<-"2"

allcurves.df$ID[allcurves.df$Tree=="4" & allcurves.df$Measure=="ACi"]<-"3"

allcurves.df$ID[allcurves.df$Tree=="5" & allcurves.df$Measure=="ACi"]<-"4"

allcurves.df$ID[allcurves.df$Tree=="9" & allcurves.df$Measure=="ACi"]<-"5"

allcurves.df$ID[allcurves.df$Tree=="11" & allcurves.df$Measure=="ACi"]<-"6"

allcurves.df$ID[allcurves.df$Tree=="12" & allcurves.df$Measure=="ACi"]<-"7"

allcurves.df$ID[allcurves.df$Tree=="13" & allcurves.df$Measure=="ACi"]<-"8"

allcurves.df$ID[allcurves.df$Tree=="14" & allcurves.df$Measure=="ACi"]<-"9"

allcurves.df$ID[allcurves.df$Tree=="15" & allcurves.df$Measure=="ACi"]<-"10"

allcurves.df$ID[allcurves.df$Tree=="16" & allcurves.df$Measure=="ACi"]<-"11"

allcurves.df$ID[allcurves.df$Tree=="17" & allcurves.df$Measure=="ACi"]<-"12"

allcurves.df$ID[allcurves.df$Tree=="18" & allcurves.df$Measure=="ACi"]<-"13"

allcurves.df$ID[allcurves.df$Tree=="19" & allcurves.df$Measure=="ACi"]<-"14"

allcurves.df$ID[allcurves.df$Tree=="20" & allcurves.df$Measure=="ACi"]<-"15"

allcurves.df$ID[allcurves.df$Tree=="21" & allcurves.df$Measure=="ACi"]<-"16"

allcurves.df$ID[allcurves.df$Tree=="22" & allcurves.df$Measure=="ACi"]<-"17"

allcurves.df$ID[allcurves.df$Tree=="23" & allcurves.df$Measure=="ACi"]<-"18"

allcurves.df$ID[allcurves.df$Tree=="24" & allcurves.df$Measure=="ACi"]<-"19"

allcurves.df$ID[allcurves.df$Tree=="2" & allcurves.df$Measure=="RACiR"]<-"21"

allcurves.df$ID[allcurves.df$Tree=="3" & allcurves.df$Measure=="RACiR"]<-"22"

allcurves.df$ID[allcurves.df$Tree=="4" & allcurves.df$Measure=="RACiR"]<-"23"

allcurves.df$ID[allcurves.df$Tree=="5" & allcurves.df$Measure=="RACiR"]<-"24"

allcurves.df$ID[allcurves.df$Tree=="9" & allcurves.df$Measure=="RACiR"]<-"25"

allcurves.df$ID[allcurves.df$Tree=="11" & allcurves.df$Measure=="RACiR"]<-"26"

allcurves.df$ID[allcurves.df$Tree=="12" & allcurves.df$Measure=="RACiR"]<-"27"

allcurves.df$ID[allcurves.df$Tree=="13" & allcurves.df$Measure=="RACiR"]<-"28"

allcurves.df$ID[allcurves.df$Tree=="14" & allcurves.df$Measure=="RACiR"]<-"29"

allcurves.df$ID[allcurves.df$Tree=="15" & allcurves.df$Measure=="RACiR"]<-"30"

allcurves.df$ID[allcurves.df$Tree=="16" & allcurves.df$Measure=="RACiR"]<-"31"

allcurves.df$ID[allcurves.df$Tree=="17" & allcurves.df$Measure=="RACiR"]<-"32"

allcurves.df$ID[allcurves.df$Tree=="18" & allcurves.df$Measure=="RACiR"]<-"33"

allcurves.df$ID[allcurves.df$Tree=="19" & allcurves.df$Measure=="RACiR"]<-"34"

allcurves.df$ID[allcurves.df$Tree=="20" & allcurves.df$Measure=="RACiR"]<-"35"

allcurves.df$ID[allcurves.df$Tree=="21" & allcurves.df$Measure=="RACiR"]<-"36"

allcurves.df$ID[allcurves.df$Tree=="22" & allcurves.df$Measure=="RACiR"]<-"37"

allcurves.df$ID[allcurves.df$Tree=="23" & allcurves.df$Measure=="RACiR"]<-"38"

allcurves.df$ID[allcurves.df$Tree=="24" & allcurves.df$Measure=="RACiR"]<-"39"

ACi.comp<-read.table("/Directory/Subdirectory/ACiR vs RACiR results.csv", header = TRUE, sep = ",", dec=".", fill = TRUE, check.names=FALSE)

ACi.comp$ID[ACi.comp$Tree=="2" & ACi.comp$Measure=="ACi"]<-"1"

ACi.comp$ID[ACi.comp$Tree=="3" & ACi.comp$Measure=="ACi"]<-"2"

ACi.comp$ID[ACi.comp$Tree=="4" & ACi.comp$Measure=="ACi"]<-"3"

ACi.comp$ID[ACi.comp$Tree=="5" & ACi.comp$Measure=="ACi"]<-"4"

ACi.comp$ID[ACi.comp$Tree=="9" & ACi.comp$Measure=="ACi"]<-"5"

ACi.comp$ID[ACi.comp$Tree=="11" & ACi.comp$Measure=="ACi"]<-"6"

ACi.comp$ID[ACi.comp$Tree=="12" & ACi.comp$Measure=="ACi"]<-"7"

ACi.comp$ID[ACi.comp$Tree=="13" & ACi.comp$Measure=="ACi"]<-"8"

ACi.comp$ID[ACi.comp$Tree=="14" & ACi.comp$Measure=="ACi"]<-"9"

ACi.comp$ID[ACi.comp$Tree=="15" & ACi.comp$Measure=="ACi"]<-"10"

ACi.comp$ID[ACi.comp$Tree=="16" & ACi.comp$Measure=="ACi"]<-"11"

ACi.comp$ID[ACi.comp$Tree=="17" & ACi.comp$Measure=="ACi"]<-"12"

ACi.comp$ID[ACi.comp$Tree=="18" & ACi.comp$Measure=="ACi"]<-"13"

ACi.comp$ID[ACi.comp$Tree=="19" & ACi.comp$Measure=="ACi"]<-"14"

ACi.comp$ID[ACi.comp$Tree=="20" & ACi.comp$Measure=="ACi"]<-"15"

ACi.comp$ID[ACi.comp$Tree=="21" & ACi.comp$Measure=="ACi"]<-"16"

ACi.comp$ID[ACi.comp$Tree=="22" & ACi.comp$Measure=="ACi"]<-"17"

ACi.comp$ID[ACi.comp$Tree=="23" & ACi.comp$Measure=="ACi"]<-"18"

ACi.comp$ID[ACi.comp$Tree=="24" & ACi.comp$Measure=="ACi"]<-"19"

ACi.comp$ID[ACi.comp$Tree=="2" & ACi.comp$Measure=="RACiR"]<-"21"

ACi.comp$ID[ACi.comp$Tree=="3" & ACi.comp$Measure=="RACiR"]<-"22"

ACi.comp$ID[ACi.comp$Tree=="4" & ACi.comp$Measure=="RACiR"]<-"23"

ACi.comp$ID[ACi.comp$Tree=="5" & ACi.comp$Measure=="RACiR"]<-"24"

ACi.comp$ID[ACi.comp$Tree=="9" & ACi.comp$Measure=="RACiR"]<-"25"

ACi.comp$ID[ACi.comp$Tree=="11" & ACi.comp$Measure=="RACiR"]<-"26"

ACi.comp$ID[ACi.comp$Tree=="12" & ACi.comp$Measure=="RACiR"]<-"27"

ACi.comp$ID[ACi.comp$Tree=="13" & ACi.comp$Measure=="RACiR"]<-"28"

ACi.comp$ID[ACi.comp$Tree=="14" & ACi.comp$Measure=="RACiR"]<-"29"

ACi.comp$ID[ACi.comp$Tree=="15" & ACi.comp$Measure=="RACiR"]<-"30"

ACi.comp$ID[ACi.comp$Tree=="16" & ACi.comp$Measure=="RACiR"]<-"31"

ACi.comp$ID[ACi.comp$Tree=="17" & ACi.comp$Measure=="RACiR"]<-"32"

ACi.comp$ID[ACi.comp$Tree=="18" & ACi.comp$Measure=="RACiR"]<-"33"

ACi.comp$ID[ACi.comp$Tree=="19" & ACi.comp$Measure=="RACiR"]<-"34"

ACi.comp$ID[ACi.comp$Tree=="20" & ACi.comp$Measure=="RACiR"]<-"35"

ACi.comp$ID[ACi.comp$Tree=="21" & ACi.comp$Measure=="RACiR"]<-"36"

ACi.comp$ID[ACi.comp$Tree=="22" & ACi.comp$Measure=="RACiR"]<-"37"

ACi.comp$ID[ACi.comp$Tree=="23" & ACi.comp$Measure=="RACiR"]<-"38"

ACi.comp$ID[ACi.comp$Tree=="24" & ACi.comp$Measure=="RACiR"]<-"39"

\#####

\#####

#Paired t-test

\#####

#Removing points below the 0.08 gsw threshold

t.test(Jmax ~ Measure,

data=subset(ACi.comp, Filtered!="All measurements"),

paired=TRUE,

conf.level=0.95)

t.test(Vcmax ~ Measure,

data=subset(ACi.comp, Filtered!="All measurements"),

paired=TRUE,

conf.level=0.95)

t.test(min.gs ~ Measure,

data=subset(ACi.comp, Filtered!="All measurements"),

paired=TRUE,

conf.level=0.95)

t.test(max.gs ~ Measure,

data=subset(ACi.comp, Filtered!="All measurements"),

paired=TRUE,

conf.level=0.95)

t.test(Rd ~ Measure,

data=subset(ACi.comp, Filtered!="All measurements"),

paired=TRUE,

conf.level=0.95)

t.test(Ci.trans ~ Measure,

data=subset(ACi.comp, Filtered!="All measurements"),

paired=TRUE,

conf.level=0.95)

t.test(Ci.range ~ Measure,

data=subset(ACi.comp, Filtered!="All measurements"),

paired=TRUE,

conf.level=0.95)

t.test(Ci.max ~ Measure,

data=subset(ACi.comp, Filtered!="All measurements"),

paired=TRUE,

conf.level=0.95)

t.test(Ci.min ~ Measure,

data=subset(ACi.comp, Filtered!="All measurements"),

paired=TRUE,

conf.level=0.95)

#Without removing points below the 0.08 gsw threshold

t.test(Jmax ~ Measure,

data=subset(ACi.comp, Filtered=="All measurements"),

paired=TRUE,

conf.level=0.95)

t.test(Vcmax ~ Measure,

data=subset(ACi.comp, Filtered=="All measurements"),

paired=TRUE,

conf.level=0.95)

t.test(min.gs ~ Measure,

data=subset(ACi.comp, Filtered=="All measurements"),

paired=TRUE,

conf.level=0.95)

t.test(max.gs ~ Measure,

data=subset(ACi.comp, Filtered=="All measurements"),

paired=TRUE,

conf.level=0.95)

t.test(Rd ~ Measure,

data=subset(ACi.comp, Filtered=="All measurements"),

paired=TRUE,

conf.level=0.95)

t.test(Ci.trans ~ Measure,

data=subset(ACi.comp, Filtered=="All measurements"),

paired=TRUE,

conf.level=0.95)

t.test(Ci.range ~ Measure,

data=subset(ACi.comp, Filtered=="All measurements"),

paired=TRUE,

conf.level=0.95)

t.test(Ci.max ~ Measure,

data=subset(ACi.comp, Filtered=="All measurements"),

paired=TRUE,

conf.level=0.95)

t.test(Ci.min ~ Measure,

data=subset(ACi.comp, Filtered=="All measurements"),

paired=TRUE,

conf.level=0.95)

\#####

#Plots

\#####

ci.range.p<-ggplot(subset(summary1, variable!="Tree"), aes(x=value,y=Type))+

geom_point(aes(color=variable), alpha=0.5)+

#geom_point(aes(x=Ci.min), color="purple", alpha=0.5)+

#geom_violin(fill="black", alpha=.3)+

theme_bw() + # remove grey background

xlab(expression(paste("C"[i]," (ppm)")))+

ylab(expression(paste("")))+

theme(panel.grid.major = element_blank()) +

scale_fill_manual(values=c("orange", "purple"))+

scale_color_manual(values=c("purple","orange" ))+

theme(axis.text.x = element_text(angle = 45, hjust = 1))

ci.range.p

gsw.range.p<-ggplot(subset(summary2,variable!="Tree"), aes(x=Type,y=value))+

geom_point(aes(color=variable), alpha=0.5,position = position_dodge(width = 0.5))+

#geom_point(aes(y=min), color="purple", alpha=0.5,position = position_nudge(x = 0.1))+

#geom_violin(fill="black", alpha=.3)+

ylim(0,.35)+

theme_bw() + # remove grey background

xlab(expression(paste("")))+

ylab(expression(paste("")))+

geom_hline(yintercept=0.08)+

theme(panel.grid.major = element_blank()) +

scale_fill_manual(values=c("orange", "purple"))+

scale_color_manual(values=c("orange", "purple"))+

theme(

#axis.text.x = element_text(angle = 45, hjust = 1),

axis.text.y = element_blank()

)

gsw.range.p

layout <- "

AAAA#

AAAA#

AAAA#

BBBBD

BBBBD

BBBBD

CCCC#

"

big.plot<-curve.p+ gsw.curve.p+ci.range.p+gsw.range.p+

plot_layout(design = layout,guides = 'collect')& theme(legend.position = 'top')

#plot_layout(heights = c(2.2,2,1),widths = c(2.5, 1))

setwd("/Users/civince/Dropbox (UFL)/Experiments/RACiR")

ggsave(filename = "RACiR curve plot.png", big.plot,width = 5, height = 8, dpi = 300, units = "in", device='png')

Vcmax.p<-ggplot(subset(ACi.comp), aes(x=Measure, y=Vcmax, group=Measure))+

facet_grid(.~Filtered)+

geom_boxplot()+

geom_jitter(shape=21)+

ylim(60,160)+

#geom_violin(fill="black", alpha=.3)+

theme_bw() + # remove grey background

ylab(expression(paste(italic("V") ["cmax"], " (", mu, "mol m" ^"-2", " sec"^"-1", ")" )))+

xlab(expression(paste("" )))+

theme(axis.text.x = element_blank(),

panel.grid.major = element_blank(),

panel.grid.minor = element_blank(),

legend.position="top",

text = element_text(size=12),

strip.background = element_blank(),

panel.border = element_rect(colour = "black"))

Vcmax.p

Jmax.p<-ggplot(subset(ACi.comp, Jmax<1000), aes(x=Measure, y=Jmax))+

facet_grid(.~Filtered)+

geom_boxplot()+

geom_jitter(shape=21)+

ylim(75,200)+

theme_bw() + # remove grey background

ylab(expression(paste(italic("J") ["1200"], " (", mu, "mol m" ^"-2", " sec"^"-1", ")" )))+

xlab(expression(paste("" )))+

theme(axis.text.x = element_blank(),

panel.grid.major = element_blank(),

panel.grid.minor = element_blank(),

legend.position="top",

text = element_text(size=12),

strip.background = element_blank(),

panel.border = element_rect(colour = "black"))

Jmax.p

Rd.p<-ggplot(subset(ACi.comp), aes(x=Measure, y=Rd))+

facet_grid(.~Filtered)+

geom_boxplot()+

geom_jitter(shape=21)+

ylim(-0.6,2.6)+

theme_bw() + # remove grey background

ylab(expression(paste(italic("R") ["d"], " (", mu, "mol m" ^"-2", " sec"^"-1", ")" )))+

xlab(expression(paste("" )))+

theme(axis.text.x = element_blank(),

panel.grid.major = element_blank(),

panel.grid.minor = element_blank(),

legend.position="top",

text = element_text(size=12),

strip.background = element_blank(),

panel.border = element_rect(colour = "black"))

Rd.p

p4<-ggplot(subset(ACi.comp), aes(interaction(Measure,Filtered),init.gs))+

geom_boxplot()+

#geom_violin(fill="black", alpha=.3)+

theme_bw() + # remove grey background

ylab(expression(paste("Initial gsw")))+

theme(panel.grid.major = element_blank()) +

theme(text = element_text(face='bold'),

axis.text.x = element_text(angle = 45, hjust = 1))

p4

p5<-ggplot(subset(ACi.comp), aes(interaction(Measure,Filtered),min.gs))+

geom_boxplot()+

#geom_violin(fill="black", alpha=.3)+

theme_bw() + # remove grey background

ylab(expression(paste("Minimum gsw")))+

theme(panel.grid.major = element_blank()) +

theme(text = element_text(face='bold'),

axis.text.x = element_text(angle = 45, hjust = 1))

p5

p6<-ggplot(subset(ACi.comp), aes(interaction(Measure,Filtered),mean.gs))+

geom_boxplot()+

#geom_violin(fill="black", alpha=.3)+

theme_bw() + # remove grey background

ylab(expression(paste("Mean gsw")))+

theme(panel.grid.major = element_blank()) +

theme(text = element_text(face='bold'),

axis.text.x = element_text(angle = 45, hjust = 1))

p6

p7<-ggplot(subset(ACi.comp), aes(interaction(Measure,Filtered),max.gs))+

geom_boxplot()+

#geom_violin(fill="black", alpha=.3)+

theme_bw() + # remove grey background

ylab(expression(paste("Maximum gsw")))+

theme(panel.grid.major = element_blank()) +

theme(text = element_text(face='bold'),

axis.text.x = element_text(angle = 45, hjust = 1))

p7

TPU.p<-ggplot(subset(ACi.comp), aes(x=Measure, y=TPU))+

facet_grid(.~Filtered)+

geom_boxplot()+

geom_jitter(shape=21)+

theme_bw() + # remove grey background

ylab(expression(paste("TPU (", mu, "mol m" ^"-2", " sec"^"-1", ")" )))+

xlab(expression(paste("" )))+

theme(axis.text.x = element_blank(),

panel.grid.major = element_blank(),

panel.grid.minor = element_blank(),

legend.position="top",

text = element_text(size=12),

strip.background = element_blank(),

panel.border = element_rect(colour = "black"))

TPU.p

p9<-ggplot(subset(ACi.comp), aes(interaction(Measure,Filtered),Ci.range))+

geom_boxplot()+

geom_jitter()+

#geom_violin(fill="black", alpha=.3)+

theme_bw() + # remove grey background

ylab(expression(paste("Magnitude of Ci range")))+

theme(panel.grid.major = element_blank()) +

theme(text = element_text(face='bold'),

axis.text.x = element_text(angle = 45, hjust = 1))

p9

Ci.trans.1<-ggplot(subset(ACi.comp, Ci.trans<1500), aes(x=Measure, y=Ci.trans))+

facet_grid(.~Filtered)+

geom_boxplot()+

geom_jitter(shape=21)+

theme_bw() + # remove grey background

ylab(expression(paste("C" [itrans1], " (", mu, "mol mol" ^"-1", ")" )))+

xlab(expression(paste("")))+

theme(panel.grid.major = element_blank()) +

theme(axis.text.x = element_text(angle = 45, hjust = 1),

panel.grid.major = element_blank(),

panel.grid.minor = element_blank(),

legend.position="top",

text = element_text(size=12),

strip.background = element_blank(),

panel.border = element_rect(colour = "black"))

Ci.trans.1

ACi.comp$Ci.trans.2<-as.numeric(as.character(ACi.comp$Ci.trans.2))

Ci.trans.2<-ggplot(subset(ACi.comp), aes(x=Measure, y=Ci.trans.2))+

facet_grid(.~Filtered)+

geom_boxplot(aes(group=Measure))+

geom_jitter(shape=21)+

#geom_violin(fill="black", alpha=.3)+

theme_bw() + # remove grey background

ylab(expression(paste("C" [itrans2], " (", mu, "mol mol" ^"-1", ")" )))+

xlab(expression(paste("")))+

theme(axis.text.x = element_text(angle = 45, hjust = 1),

panel.grid.major = element_blank(),

panel.grid.minor = element_blank(),

legend.position="top",

text = element_text(size=12),

strip.background = element_blank(),

panel.border = element_rect(colour = "black"))

Ci.trans.2

boxplots.p<-(Vcmax.p+Jmax.p)/(Rd.p+TPU.p)/(Ci.trans.1+Ci.trans.2)

setwd("/Users/civince/Dropbox (UFL)/Experiments/RACiR")

ggsave(filename = "Boxplots redo.png", boxplots.p,width = 6.5, height = 8, dpi = 300, units = "in", device='png')

p8<-ggplot(subset(ACi.comp, init.gs>0.05),aes(mean.gs,Vcmax))+geom_point(aes(color=Measure))+

theme_bw()+

theme(panel.grid.major = element_blank(),

panel.grid.minor = element_blank(),

legend.position="top",

text = element_text(size=12),

axis.text.x = element_text(angle = 45, hjust = 1),

strip.background = element_blank(),

panel.border = element_rect(colour = "black"))

p8

p9<-ggplot(subset(ACi.comp, init.gs>0.05),aes(mean.gs,Jmax))+geom_point(aes(color=Measure))+

theme_bw()+

theme(panel.grid.major = element_blank(),

panel.grid.minor = element_blank(),

legend.position="top",

text = element_text(size=12),

axis.text.x = element_text(angle = 45, hjust = 1),

strip.background = element_blank(),

panel.border = element_rect(colour = "black"))

p9

p10<-ggplot(subset(ACi.comp, init.gs>0.05),aes(mean.gs,Rd))+geom_point(aes(color=Measure))+

theme_bw()+

theme(panel.grid.major = element_blank(),

panel.grid.minor = element_blank(),

legend.position="top",

text = element_text(size=12),

axis.text.x = element_text(angle = 45, hjust = 1),

strip.background = element_blank(),

panel.border = element_rect(colour = "black"))

p10

p8/p9/p10+ plot_layout(guides = "collect")& theme(legend.position = 'top')

\#####

#Reanalyzing excluding gsw below 0.06

\#####

library(dplyr)

X<-allcurves.df %>%

mutate(threshold=ifelse(gsw>0.08,"TRUE","FALSE"),

CaCidiff= Ca-Ci) %>%

filter(threshold=="FALSE")%>%

group_by(Tree,Type,threshold)%>%

summarize(number=length(A))

allcurves.df<- allcurves.df%>%

mutate(Type=recode_factor(Type, `Steady state` = "SS",`RACiR` = "RACiR"))

summary1<-allcurves.df %>%

group_by(Type,Tree) %>%

summarise(maximum=max(Ci), minimum=min(Ci))%>%

melt()

summary2<-allcurves.df %>%

group_by(Type,Tree) %>%

summarise(minimum=min(gsw), maximum=max(gsw))%>%

melt()

\#####

#4

\#####

acifit <- fitaci(data=subset(allcurves.adj.df, Tree=="4"&Type=="RACiR"), varnames = list(ALEAF = "A", Tleaf = "TleafCnd",

Ci = "Ci",

PPFD = "Qin"),

fitmethod = "bilinear" ,

Tcorrect = FALSE,

fitTPU = TRUE)

acifit$Ci_transition

acifit$Ci_transition2

plot(acifit)

coef(acifit)

R.coef.<-as.data.frame(coef(acifit))

\#####

#5

\#####

acifit <- fitaci(data=subset(allcurves.adj.df, Tree=="5"&Type=="RACiR"), varnames = list(ALEAF = "A", Tleaf = "TleafCnd",

Ci = "Ci",

PPFD = "Qin"),

fitmethod = "bilinear" ,

Tcorrect = FALSE,

fitTPU = TRUE)

acifit$Ci_transition

acifit$Ci_transition2

plot(acifit)

coef(acifit)

R.coef.<-as.data.frame(coef(acifit))

\#####

#14

\#####

acifit <- fitaci(data=subset(allcurves.adj.df, Tree=="14"&Type=="RACiR"), varnames = list(ALEAF = "A", Tleaf = "TleafCnd",

Ci = "Ci",

PPFD = "Qin"),

fitmethod = "bilinear" ,

Tcorrect = FALSE,

fitTPU = TRUE)

acifit$Ci_transition

acifit$Ci_transition2

plot(acifit)

coef(acifit)

R.coef.<-as.data.frame(coef(acifit))

\#####

#ACi

\####

\#####

#2

\#####

acifit <- fitaci(data=subset(allcurves.adj.df, Tree=="2"&Type=="Aci"), varnames = list(ALEAF = "A", Tleaf = "TleafCnd",

Ci = "Ci",

PPFD = "Qin"),

fitmethod = "bilinear" ,

Tcorrect = FALSE,

fitTPU = TRUE)

acifit$Ci_transition

acifit$Ci_transition2

plot(acifit)

coef(acifit)

R.coef.<-as.data.frame(coef(acifit))

\#####

#4

\#####

acifit <- fitaci(data=subset(allcurves.adj.df, Tree=="4"&Type=="Aci"), varnames = list(ALEAF = "A", Tleaf = "TleafCnd",

Ci = "Ci",

PPFD = "Qin"),

fitmethod = "bilinear" ,

Tcorrect = FALSE,

fitTPU = TRUE)

acifit$Ci_transition

acifit$Ci_transition2

plot(acifit)

coef(acifit)

R.coef.<-as.data.frame(coef(acifit))

\#####

#13

\#####

acifit <- fitaci(data=subset(allcurves.adj.df, Tree=="13"&Type=="Aci"), varnames = list(ALEAF = "A", Tleaf = "TleafCnd",

Ci = "Ci",

PPFD = "Qin"),

fitmethod = "bilinear" ,

Tcorrect = FALSE,

fitTPU = TRUE)

acifit$Ci_transition

acifit$Ci_transition2

plot(acifit)

coef(acifit)

R.coef.<-as.data.frame(coef(acifit))

\#####

#14

\#####

acifit <- fitaci(data=subset(allcurves.adj.df, Tree=="14"&Type=="Aci"), varnames = list(ALEAF = "A", Tleaf = "TleafCnd",

Ci = "Ci",

PPFD = "Qin"),

fitmethod = "bilinear" ,

Tcorrect = FALSE,

fitTPU = TRUE)

acifit$Ci_transition

acifit$Ci_transition2

plot(acifit)

coef(acifit)

R.coef.<-as.data.frame(coef(acifit))

\#####

#15

\#####

acifit <- fitaci(data=subset(allcurves.adj.df, Tree=="15"&Type=="Aci"), varnames = list(ALEAF = "A", Tleaf = "TleafCnd",

Ci = "Ci",

PPFD = "Qin"),

fitmethod = "bilinear" ,

Tcorrect = FALSE,

fitTPU = TRUE)

acifit$Ci_transition

acifit$Ci_transition2

plot(acifit)

coef(acifit)

R.coef.<-as.data.frame(coef(acifit))

\#####

#16

\#####

acifit <- fitaci(data=subset(allcurves.adj.df, Tree=="16"&Type=="Aci"), varnames = list(ALEAF = "A", Tleaf = "TleafCnd",

Ci = "Ci",

PPFD = "Qin"),

fitmethod = "bilinear" ,

Tcorrect = FALSE,

fitTPU = TRUE)

acifit$Ci_transition

acifit$Ci_transition2

plot(acifit)

coef(acifit)

R.coef.<-as.data.frame(coef(acifit))

\#####

#17

\#####

acifit <- fitaci(data=subset(allcurves.adj.df, Tree=="17"&Type=="Aci"), varnames = list(ALEAF = "A", Tleaf = "TleafCnd",

Ci = "Ci",

PPFD = "Qin"),

fitmethod = "bilinear" ,

Tcorrect = FALSE,

fitTPU = TRUE)

acifit$Ci_transition

acifit$Ci_transition2

plot(acifit)

coef(acifit)

R.coef.<-as.data.frame(coef(acifit))

\#####

#18

\#####

acifit <- fitaci(data=subset(allcurves.adj.df, Tree=="18"&Type=="Aci"), varnames = list(ALEAF = "A", Tleaf = "TleafCnd",

Ci = "Ci",

PPFD = "Qin"),

fitmethod = "bilinear" ,

Tcorrect = FALSE,

fitTPU = TRUE)

acifit$Ci_transition

acifit$Ci_transition2

plot(acifit)

coef(acifit)

R.coef.<-as.data.frame(coef(acifit))

\#####

#19

\#####

acifit <- fitaci(data=subset(allcurves.adj.df, Tree=="19"&Type=="Aci"), varnames = list(ALEAF = "A", Tleaf = "TleafCnd",

Ci = "Ci",

PPFD = "Qin"),

fitmethod = "bilinear" ,

Tcorrect = FALSE,

fitTPU = TRUE)

acifit$Ci_transition

acifit$Ci_transition2

plot(acifit)

coef(acifit)

R.coef.<-as.data.frame(coef(acifit))

\#####

#20

\#####

acifit <- fitaci(data=subset(allcurves.adj.df, Tree=="20"&Type=="Aci"), varnames = list(ALEAF = "A", Tleaf = "TleafCnd",

Ci = "Ci",

PPFD = "Qin"),

fitmethod = "bilinear" ,

Tcorrect = FALSE,

fitTPU = TRUE)

acifit$Ci_transition

acifit$Ci_transition2

plot(acifit)

coef(acifit)

R.coef.<-as.data.frame(coef(acifit))

\#####

#22

\#####

acifit <- fitaci(data=subset(allcurves.adj.df, Tree=="22"&Type=="Aci"), varnames = list(ALEAF = "A", Tleaf = "TleafCnd",

Ci = "Ci",

PPFD = "Qin"),

fitmethod = "bilinear" ,

Tcorrect = FALSE,

fitTPU = TRUE)

acifit$Ci_transition

acifit$Ci_transition2

plot(acifit)

coef(acifit)

R.coef.<-as.data.frame(coef(acifit))

\#####

#23

\#####

acifit <- fitaci(data=subset(allcurves.adj.df, Tree=="23"&Type=="Aci"), varnames = list(ALEAF = "A", Tleaf = "TleafCnd",

Ci = "Ci",

PPFD = "Qin"),

fitmethod = "bilinear" ,

Tcorrect = FALSE,

fitTPU = TRUE)

acifit$Ci_transition

acifit$Ci_transition2

plot(acifit)

coef(acifit)

R.coef.<-as.data.frame(coef(acifit))

\#####

#24

\#####

acifit <- fitaci(data=subset(allcurves.adj.df, Tree=="24"&Type=="Aci"), varnames = list(ALEAF = "A", Tleaf = "TleafCnd",

Ci = "Ci",

PPFD = "Qin"),

fitmethod = "bilinear" ,

Tcorrect = FALSE,

fitTPU = TRUE)

acifit$Ci_transition

acifit$Ci_transition2

plot(acifit)

coef(acifit)

R.coef.<-as.data.frame(coef(acifit))
